# Methodological pitfalls in plant pangenome gene family identification may lead to biased evolutionary inferences

**DOI:** 10.64898/2026.05.15.725319

**Authors:** Shuotong Liu, Wei Zhang, Pei Yu

## Abstract

Pangenome-level gene family identification often applies sequence similarity clustering without phylogenetic or synteny information, which risks biologically misleading evolutionary inferences. Using five transcription factor families (bHLH, MYB, NAC, WRKY, MADS-box) across 401 rice pangenome accessions, we compared clustering strategies: OrthoFinder alone, cd-hit alone, MMseqs2 alone, and OrthoFinder-informed refinement by cd-hit or MMseqs2. Methods solely based on sequence similarity merged distinct orthogroups and generated fewer orthogroups than approaches incorporating graph-based orthology. Conflicting cluster assignments, measured against OrthoFinder, varied strongly among families, from approximately 14% in MADS-box to approximately 57% in MYB, and were associated with protein length differences. Core, shell, and cloud gene classifications shifted substantially depending on the method, especially in MYB, NAC, and WRKY families. Critically, Ka/Ks distributions for core genes were highly method-sensitive, with orthology-aware methods yielding more convergent and less variable estimates of selective pressure, whereas noncore gene estimates remained robust. These findings demonstrate that neglecting graph-based orthogroup inference inflates methodological artifacts. We recommend a two-step strategy: initial graph-based orthogroup delineation followed by sequence similarity refinement to balance evolutionary accuracy and resolution in pangenome-scale gene family studies.

## Dear Editor

The research paradigm of gene family identification and analysis has been expanded to the pangenome level based on multiple reference genomes (Tong et al., 2025). However, after the multi-reference identification was completed, there was no unified standard for the inferring of orthologous gene groups. Some studies used cd-hit/MMseqs2 or OrthoFinder (Li and Godzik, 2006; Steinegger and Söding, 2017; Emms and Kelly, 2019) alone to obtain the orthologous group in one gene family, which was insufficiently robust and did not incorporate phylogenetic and synteny information, are intrinsically problematic (Huang et al., 2025; Liu and Yu, 2025; Edwards, 2026).

Through the pangenome-wide identification of five common transcription factor (TF) families in 401 rice pangenome accessions (Shang et al., 2022; Guo et al., 2025), we propose to consider both graph-based insight and sequence similarity when clustering orthologous gene groups, thereby avoiding biologically misleading conclusions by methodological flaws.

We compared clustering methods based purely on sequence similarity (cd-hit, MMseqs2) with those integrating graph-based clustering (via Markov Cluster Algorithm (MCL), OrthoFinder-based strategies) (Fig. 1A). We have defined five cluster analysis methods. Method 1 uses OrthoFinder alone. Methods 2 and 3 use cd-hit or MMseqs2 alone. Method 4 and 5 introduce cd-hit or MMseqs2 on the basis of using OrthoFindner.

**Figure 1.**
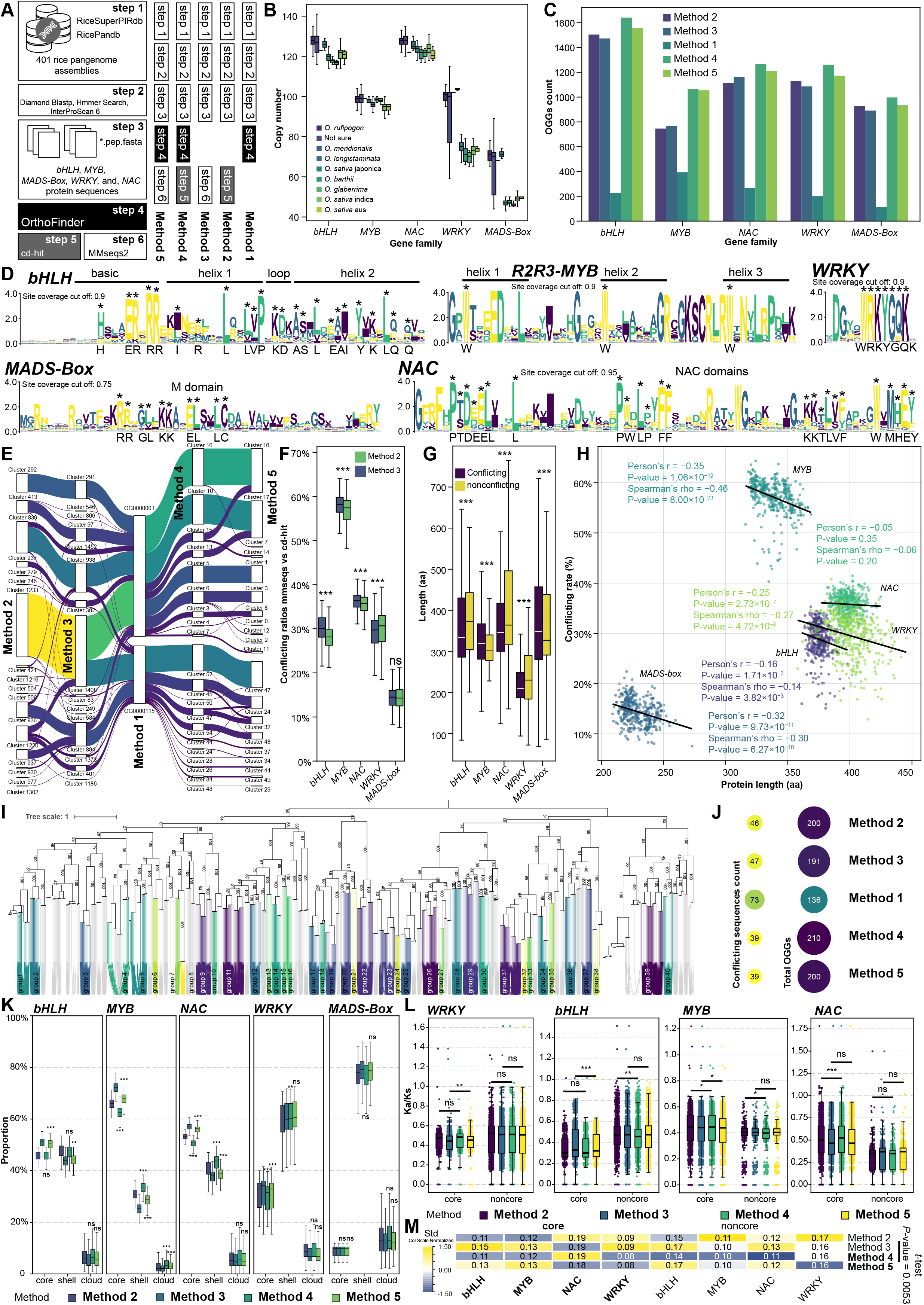
Five plant transcription factor (TF) families between five different methods to infer OGGs in rice pangenome. **(A)** Schematic workflow of this study, illustrating the key procedural differences among the five clustering methods evaluated. **(B)** Copy number of the *bHLH, MYB, NAC, WRKY* and *MADS-Box* family in rice pangenome with species differentiated. **(C)** The comparison of the numbers of OGGs in five different methods. **(D)** Sequence alignment and conserved motif validation of five TF families. **(E)** A Sankey diagram illustrating the mis-clustering of orthogroups caused by sequence-similarity-based clustering methods. **(F)** Comparison of conflicting ratios between two clustering methods (MMseqs2 vs cd-hit) across five plant TF families. Statistical significance was determined using Mann-Whitney *U* tests. “***” indicates a *P* < 0.001, while “ns” denotes not significant. **(G)** Protein length distribution in each TF family. Statistical significance was determined same as Fig. 1F. The conflicting classification indicates sequences that were assigned to different orthogroups by similarity-based methods (Methods 2/3) relative to the OrthoFinder reference grouping. **(H)** Relationship between protein length (aa) and clustering conflicting rate across different TF families. Each scatter point represents a distinct assembly from the rice pangenome. Data points for different TF families are distinguished and fitted with separate trend lines. The strength and significance of the correlations were assessed using both Pearson’s *r* and Spearman’s *ρ*, with corresponding *P* calculated for each family. **(I)** Comparative analysis of collinearity and phylogenetic relationships of *bHLH* in five *O. sativa* japonica accessions (NIP-T2T, Nip_hifi, NH001, HP436, and, GP117). **(J)** The heatmap of conflicting and total OGGs in different methods. The conflicting OGGs means different clusters in collinear networks which are assigned in the same OGGs in the five methods. **(K)** Distribution patterns of core, shell, and cloud gene categories across five TF families identified by different clustering methods. For each family, the proportions of genes classified as core (conserved in ≥90% lines), shell (>10%), and cloud (accession-specific, ≤10%) are shown. Statistical comparisons of the distribution patterns between methods were performed using the Mann-Whitney *U* test. Significance levels are indicated as “***” for *P* < 0.001, “**” for *P* < 0.01, “*” for *P* < 0.05, and “ns” for not significant. **(L)** Distribution of Ka/Ks ratios for core and noncore gene pairs across four TF families inferred by different clustering methods. The Ka/Ks ratio (nonsynonymous substitution rate per nonsynonymous site divided by synonymous substitution rate per synonymous site). “core” refers to gene pairs where both genes are classified as core (conserved in ≥90% lines), while “noncore” refers to pairs where neither gene is classified as core. **(M)** Heatmap displaying the normalized standard deviation (Std) of Ka/Ks ratios for “core” and “noncore” gene pairs across four TF families under different clustering methods. Values were normalized by column to enable cross-family and cross-category comparison. Paired *t*-tests were performed using raw data without normalizing to evaluate the effect across all families and categories.

Among the five TF gene families examined, *NAC* and *bHLH* exhibited the highest copy numbers, whereas *MADS-Box* showed the lowest (Fig. 1B). The five clustering methods yielded markedly different numbers of orthologous groups. Clustering using OrthoFinder alone (Method 1) produced the fewest orthogroups across all families (Fig.1C). Although cd-hit (Method 2) and MMseqs2 (Method 3) are algorithmically distinct, the number of clusters generated by these two methods was comparable. In contrast, clustering informed by OrthoFinder (Methods 4 and 5), which integrates both sequence similarity and graph-based relationships, resulted in substantially more clusters than either cd-hit or MMseqs2 alone (Fig. 1C).

To ensure the accuracy of subsequent validation, we aligned and trimmed the sequences of the five TF families and compared them with results from previous studies, further confirming the reliability of our current identifications. Taking the *bHLH* family as an example, the conserved motifs—basic region: H, E, R, R, R; helix 1: I, R, L, L, V, P; loop: K, D; helix 2: A, S, L, E, A, I, Y, K, L, Q, Q, and the conserved sites of *WRKY*: W, R, K, Y, G, Q, K—almost perfectly matched the previously reported consensus sequences (Feller et al., 2011; Fig. 1D).

In the *bHLH* family, cluster 1408, generated by Method 3, merged members from distinct orthogroups (OG0000001 and OG0000115) (Fig.1E). Although this represents a specific case, it illustrates a broader trend: without the constraints of graph-based orthology inference, switching between similarity-based algorithms fails to resolve fundamental misclassifications. Moreover, notable variation in conflicting ratios was observed across different TF families (Fig. 1F). Conflicting ratios varied substantially, ranging from ∼14% in *MADS-box* to ∼57% in *MYB*, with method 3 (MMseqs2) generally yielding slightly higher rates than method 2 (cd-hit) (Fig. 1F), suggesting that sequence divergence among families may underlie the inconsistent clustering performance between methods. It should be pointed out that “conflicting” refers to sequences that were assigned to a different orthogroup by the tested method compared to the reference orthogroup assignment in OrthoFinder. This reference was chosen for the purpose of method comparison, rather than treating OrthoFinder as the absolute truth.

To investigate factors contributing to clustering errors, we compared sequence lengths between conflicting and nonconflicting clustered sequences for each family (Fig. 1G). Highly significant differences were observed across all five families, although the direction varied. In *bHLH, NAC* and WRKY families, conflicting clustered sequences were significantly longer than nonconflicting ones. Conversely, in *MYB* and *MADS-box* families, clustered sequences were significantly shorter. Sequence length is strongly associated with clustering conflicts, but whether sequences leading to conflicts tend to be longer or shorter, the current analytical methods have yet to reach a definite conclusion.

We next examined whether this sequence-level association extends to the conflicting rate level (Fig. 1H). For each family across 401 rice pangenome assemblies, we calculated average protein length and conflicting rate per assembly and analyzed their correlation. A general negative trend between protein length and conflicting rate was observed across most families, with *MYB* showing the strongest correlation and *NAC* showing no significant correlation (Fig. 1H). These findings indicate that protein length is a consistent factor influencing clustering accuracy across most families, with longer sequences generally associated with lower conflicting rates at the assembly level (Fig. 1H).

Notablely, these two analyses operate at different scales—sequence-level differences within each family (Fig. 1G) versus assembly-level correlations across families (Fig. 1H) —and their findings are complementary rather than contradictory.

Further analysis using partial genome assemblies and syntenic relationships confirmed that the collinearity networks generated by MCScanX (40 main groups in total) were largely consistent with phylogenetic trees (Fig. 1I). Under comparable total numbers of OGGs, Methods 4/5 yielded substantially fewer conflicting orthogroups than Methods 2/3, further validating the benefit of incorporating OrthoFinder (Fig. 1J).

The classification of core, shell, and cloud genes was highly sensitive to clustering method (Fig. 1K). While MADS-box showed robust classifications across methods, the *WRKY* family, alongside *bHLH, MYB*, and, *NAC*, displayed significant discrepancies, particularly when comparing methods with (Methods 4/5) and without (Methods 2/3) OrthoFinder preprocessing. The effect was family-dependent: for *MYB, NAC* and *WRKY*, incorporating OrthoFinder significantly altered core/shell proportions, whereas for *bHLH* and *MADS-box*, patterns were more stable. Cloud genes were largely unaffected. Importantly, integrating graph-based information via OrthoFinder preprocessing (Methods 4/5) substantially alters the inferred proportions of core and shell genes compared to similarity-only methods (Methods 2/3), underscoring the necessity of orthology-aware approaches for accurate pangenome structure inference (Fig. 1K).

In *WRKY* and *bHLH* families: core gene pairs are under significantly stronger purifying selection than “noncore” pairs (Fig. 1L). This supports the biological intuition that genes shared by every individual in a pangenome (core) perform essential and functionally constrained roles. However, the result is the opposite in the *NAC* and MYB family. The specific reasons for this still remain to be explored.

The choice of clustering method substantially impacted Ka/Ks distributions, particularly for core genes (Fig. 1L). In *bHLH* and *NAC*, sequence-similarity-only methods (Methods 2/3) produced significantly different Ka/Ks distributions compared to their phylogenetically-informed counterparts (Methods 4/5). The two orthology-aware methods (Methods 4/5) yielded highly similar distributions to each other in *NAC* and *MYB* core sets, suggesting that incorporating orthology provides more convergent and robust evolutionary estimates (Fig. 1L).

In contrast to core genes, the Ka/Ks distributions for “noncore” genes were largely consistent across all four clustering methods (Fig. 1L). This suggests that while the precise delineation of core orthogroups is method-sensitive, the broader group of noncore, more flexible genes is less affected by clustering strategy when estimating selective pressure. Furthermore, orthology-aware methods not only affected the central tendency but also significantly reduced the variability of Ka/Ks estimates (*P* = 0.0053), underscoring their robustness (Fig. 1M).

In summary, this study demonstrates that pangenome-scale gene family analysis requires a hybrid methodological approach. Adjusting inflation alone is often insufficient; excessively high values may lead to artificial over-splitting, while low values may fail to separate distinct subgroups. To overcome this, we recommend a two-step strategy: employing OrthoFinder for initial, stable orthogroup delineation, and subsequently applying cd-hit/MMseqs2 to achieve high-resolution clustering within those groups. The Methods 4 and 5 address a critical trade-off: while pure similarity-based methods risk merging independent evolutionary lineages, relying solely on OrthoFinder parameters can lead to arbitrary over-splitting that lacks biological granularity. By integrating graph-based grouping (Emms and Kelly, 2015; Emms and Kelly, 2019) with similarity-based refinement, we demonstrate a superior balance between evolutionary accuracy and classification resolution. This graph-based-first, similarity-second workflow (Methods 4 and 5) preserves the global evolutionary context while enabling the precise identification of recently expanded subfamilies. While this approach approximates evolutionary relationships more effectively than pairwise alignments, it remains a heuristic proxy that lacks the explicit resolution of full phylogenetic tree reconstruction. We recommend that when extending these findings to other species or different types of gene families, the applicability of the methods should be evaluated on a case-by-case basis, and preference should be given to orthology inference strategies that integrate multiple lines of evidence (sequence, phylogeny, synteny).

## Author contributions

Conceptualization, P.Y. and S.L.; Data curation, S.L.; Funding acquisition, W.Z. and P.Y.; Resources, S.L.; Bioinformatic Analysis, S.L.; Supervision, W.Z. and P.Y.; Validation, S.L.; Writing—original draft, S.L.; Writing—review & editing, P.Y. and S.L. All authors have read and agreed to the published version of the manuscript.

## Data availability

All data supporting the findings of this study is available in the article and the supplemental information files.

## Acknowledgments

Appreciation is extended to: Xiaoyi Lyu (SDU-ANU Joint Science College, Shandong University); Yueduo Wang (Radiation Biology Center, Kyoto University); Shenglong Kan (Marine College, Shandong University).

