## Supplemental information for "Methodological pitfalls in plant pangenome gene family identification may lead to biased evolutionary inferences"

Shuotong Liu^1^, Feifei Yuan ^2^, Youlong Ke^2^, Shenglong Ding^2^, Ying Zhang^2^, Qian Liu^2^, Zhen Zhang^2^, Yun Zhou^2^, Wei Zhang^3^*, Pei Yu^1^*

^1^ SDU-ANU Joint Science College, Shandong University, Weihai 264209, China

^2^ State Key Laboratory of Crop Stress Adaptation and Improvement, School of Life Sciences, Henan University, Kaifeng 475004, China

^3^ Marine College, Shandong University, Weihai 264209, China

*Corresponding authors:

Wei Zhang,,

Pei Yu,

**This PDF file includes:**

Materials and Methods

Supplemental References

Materials and methods

**Acquisition of genomic data**

The genomic data of 401 rice cultivars was downloaded from two online databases (RicePandb: <http://ricepandb.ncgr.ac.cn/home>; RiceSuperPIRdb: <http://www.ricesuperpir.com/>) (Shang et al., 2022; Guo et al., 2025). All genome files were formatted and standardized with the use of TBtools v2.458 (Chen et al., 2020; Chen et al., 2023) to ensure compatibility for subsequent analysis. More information was in Table S1.

**Identification of five plant TFs**

The protein sequences of *bHLH* gene family in *Arabidopsis thaliana* were used for Blastp in DIAMOND to accelerate the computing process. And the hidden Markov model (HMM) profile of the *bHLH* domain (PF00010) was download from InterPro (<https://www.ebi.ac.uk/interpro/>) (Blum et al., 2025). E-value was 1e−5 for both BLASTP and hmmscan from the HMMER package. Then InterProScan v6 (Jones et al., 2014; Blum et al., 2025) was used to check the domain of identification results. Proteins with more than two *bHLH* domains were deleted. The identification methods of the other 4 plant TFs were same as methods of the *bHLH* gene family (*MYB*: PF00249; *WRKY*: PF03106; *NAC*: PF02365; *MADS-Box*: PF00319 or PF01486). Regarding the complexity of the *MADS-Box* domain, we did not use a reference sequence to perform BLASTP analysis. More information was in Table S2.

**Phylogenetic tree construction**

MAFFT v7.525 was used to align the sequence in “auto” mode (Katoh and Standley, 2013). And ClipKIT v2.12.0 was used to delete non-conserved amino acid in “kpic” and “gappy” model (Steenwyk et al., 2020). The unrooted maximum-like-lihood tree was constructed using IQ-TREE v1.6.12 (Nguyen et al., 2015) with 1000 UltraFast Bootstrap evaluation of node support (Hoang et al., 2018), and ModelFinder to prefer the best alternative model (‘VT+F+R6’).

**Inference of orthologous gene groups (OGGs)**

5 methods were used in this study. In the first method, OrthoFinder (Emms and Kelly, 2015; Emms and Kelly, 2019) was used. And cd-hit (Li and Godzik, 2006) or MMSeqs2 (Steinegger and Söding, 2017) with a cutoff of 95% identity and 90% alignment (Tong et al., 2025) were applied in the 2^nd^ and 3^rd^ method. In the 4^th^ and 5^th^ method, both OrthoFinder and cd-hit or MMSeqs2 were used. Moreover, cd-hit or MMseqs2 were used after obtaining relatively rough results of gene family clustering through OrthoFinder. And the standard for cd-hit or MMSeqs2 were same as the 2^nd^ and 3^rd^ method. More information was in Table S3.

**Analysis of gene duplication and synteny**

DIAMOND (Buchfink et al., 2015) and MCScanX (Wang et al., 2012) was used to analyze gene duplication events and synteny. The protein sequences for each of the 401 pan-genome accessions were used for BLASTP analyses (E-value threshold 1e−5). Only the top 10 BLASTP hits were retained following the tool’s instructions. Both self-collinearity and collinearity among varieties were analyzed. Simultaneously, the automated identification and classification statistics of segmental and tandem duplication events were implemented through duplicate gene classifier program in MCScanX (Wang et al., 2012). More information was in Table S4.

**Analysis of natural selection**

Homologous gene pairs in collinearity analysis were analyzed using the Ka, Ks, Ka/Ks Calculator plugin in TBtools v2.458 (Simple Ka/Ks Calculator) to calculate the natural selection (Chen et al., 2020; Chen et al., 2023). Gene pairs with average sequence coverage below 70% were deleted from the Ks/Ka analysis to ensure the robustness. More information was in Table S5.

**Data plotting and statistical analysis**

Heatmaps were generated using TBtools v2.458 (Chen et al., 2020; Chen et al., 2023), and network topologies were constructed and visualized in Gephi v0.10.1 (Bastian et al., 2009). The auxiliary charts were implemented by Python. The key mathematical verification was completed through Python’s SciPy scientific computing library (Virtanen et al., 2020a; Virtanen et al., 2020b). For phylogenetic trees, iTOL was used for visualization (Letunic and Bork, 2024).
